# Diverse effects of fluorescent labels on alpha-synuclein condensate formation during liquid-liquid phase separation

**DOI:** 10.1101/2024.07.05.602219

**Authors:** Mantas Ziaunys, Darius Sulskis, Dominykas Veiveris, Andrius Sakalauskas, Kamile Mikalauskaite, Vytautas Smirnovas

## Abstract

Liquid-liquid phase separation is an emerging field of study, dedicated to understanding the mechanism and role of biomolecule assembly into membraneless organelles. One of the main methods employed in studying protein and nucleic acid droplet formation is fluorescence microscopy. Despite functioning as an excellent tool for monitoring biomolecule condensation, a few recent reports have presented possible drawbacks of using fluorescently labeled particles. It was observed that fluorescent tags could alter the process of protein liquid-liquid phase separation and even promote their aggregation. In this study, we examined the influence of three different protein labels on alpha-synuclein phase separation *in vitro* and determined that the changes in droplet formation were related to both the type, as well as concentration of the fluorescently tagged alpha-synuclein. Both protein-based labels (mCherry and eGFP) induced the formation of significantly larger droplets, while fluorescein-tagged alpha-synuclein generated an abundance of small condensates. The study also revealed that alpha-synuclein with protein-based labels could self-associate at much lower concentrations than its untagged counterpart, forming either large droplets or protein aggregates.

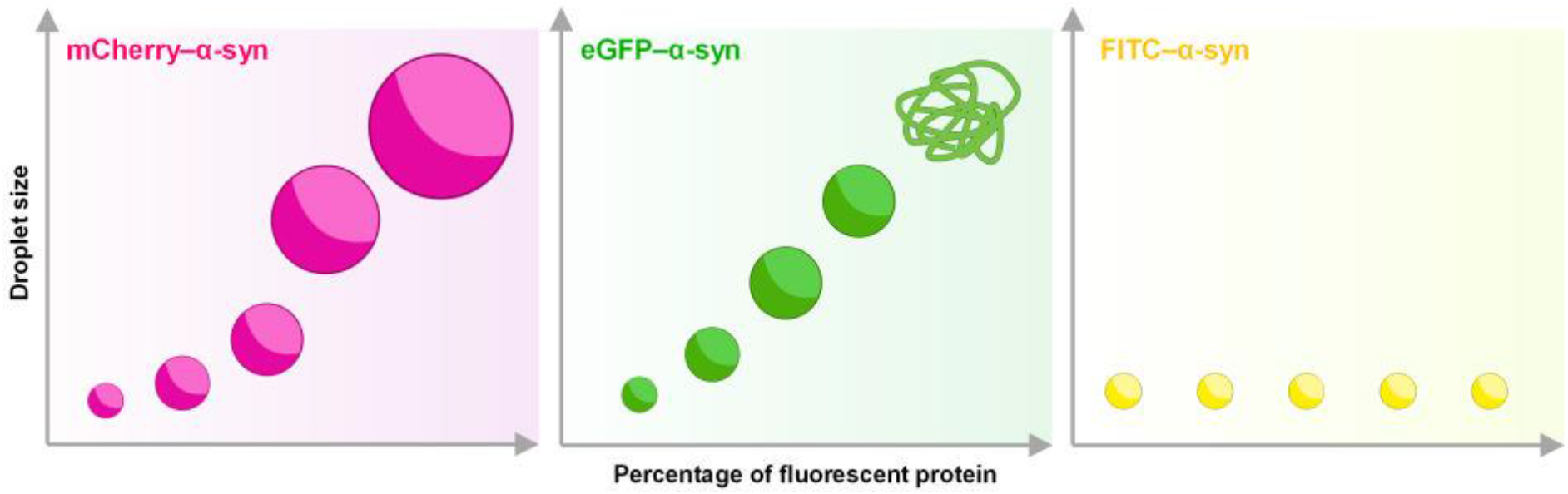

## Introduction

Liquid-liquid phase separation (LLPS) of proteins and nucleic acids is a rapidly expanding field of study, which aims at deciphering the mechanism of biomolecular condensate formation and their role in various cellular functions ^1,2^. During the last decade, this phenomenon has gained attraction as a result of several important discoveries. LLPS has been shown to be involved in the generation of promyelocytic leukemia protein bodies, stress granules, nuclear speckles and over twenty other types of condensates ^1,3–5^. In addition to a large number of regulatory roles, aberrant LLPS ^6^ has also been implicated in the onset of several widespread disorders, such as Alzheimer’s ^7^ or Parkinson’s ^8^ disease and various forms of cancer^3,4,9,10^.

For protein LLPS studies, fluorescent markers are often used to visualize and track the formation of condensates via microscopy ^11^. During these procedures, it is common that the samples contain a small fraction of particles modified with various protein-based fluorescent tags, with some of the most common ones being green (GFP, eGFP ^12^), red (RFP ^13^, mCherry ^14^) and yellow (YFP ^15^) fluorescent proteins. Alternatively, the sample can be chemically modified with a number of small fluorescent probes, including fluorescein ^8^, rhodamine ^16^ or AlexaFluor variants ^17^. However, despite these widely used techniques, little is known in regard to how such additives may influence the process of LLPS ^18^.

A few recent studies have suggested that fluorescent labels may play a critical role in protein condensate formation. In one case, it was observed that the addition of a red fluorescent protein (RFP) tag to huntingtin exon-1 strongly promoted its LLPS and even altered the protein’s amyloid aggregation pathway ^19^. Experiments conducted on alpha-crystallin variants displayed a correlation between protein tagging with eGFP and their ability to form droplets ^20^. The ability of GFP to modulate protein phase separation was also shown in a study conducted on heterochromatin protein HP1α ^21^. Another report demonstrated that different fluorescent tags (mKate2 and mTagBFP2) can alter the dynamics of P granule phase separation ^22^. These results imply that the addition of such modified proteins into the sample can alter the outcome of the experiment by distorting the LLPS process.

In addition to the large variety of fluorescent protein labels and their possible role in LLPS, there is also no consensus on the optimal concentration of the tagged particles. A review of recent literature shows that during fluorescence microscopy studies of LLPS, the amount of labeled molecules can range from as low as 0.2% - 1.0% ^15,23^ of the total concentration, to conditions where the entire sample consists of only tagged particles ^14,24^. There is also a massive variance in the total concentration of tagged proteins used, ranging from 1 µM ^25^ to 500 µM ^26^. Taking into account the large number of studies conducted with the use of fluorescence microscopy techniques, the recent reports of label-specific modulation of condensate formation, as well as the variety of concentrations and types of protein tags, it is likely that the field of LLPS has been significantly impacted by their droplet-modulating properties.

In this work, alpha-synuclein (α-syn) was selected as the basis for the investigation of different fluorescent tags and their influence on protein LLPS. α-syn is an intrinsically disordered protein ^27^, whose aggregation into amyloid fibrils is linked with the onset and progression of Parkinson’s disease ^28^. It has also been the subject of numerous LLPS studies and has been shown to easily form protein condensates *in vitro* ^29,30^, making it a perfect candidate for this examination. An analysis was conducted on how different concentrations of modified α-syn variants (tagged with eGFP, mCherry and fluorescein isothiocyanate) modulated the protein’s condensate formation process. It was observed that both mCherry and eGFP had a concentration-dependent effect and induced the assembly of significantly enlarged droplets. In addition, the eGFP tag also promoted rapid α-syn assembly into amorphous protein aggregates. Conversely, higher concentrations of fluorescein-tagged α-syn displayed large quantities of tiny protein assemblies, with no notable appearance of larger droplets. These results highlight the drastic effect that fluorescent tags have on protein LLPS and how caution should be exercised when selecting their type and concentration.

## Materials and Methods

### Protein purification

The purification of α-syn and its tagged forms (eGFP-α-syn and mCherry-α-syn) was done similarly as in previous work ^31^. In short, the expression of human α-synuclein was done in *E*.*Coli* cells (BL-21Star™ (DE3)) by culturing them overnight in the auto-induction medium (ZYM-5052) containing 100 µg/mL ampicillin at 37 °C. Once the synthesis was completed, the cells were collected by centrifuging the medium at 4 000 x g for 20 minutes at 4 °C. The cell pellet was homogenized and lysed in a buffer solution containing 20 mM Tris-HCl, 0.5 M NaCl, 1 mM EDTA and 1 mM PMSF (pH 8.0). The supernatant was subjected to a 70 °C water bath for thermal denaturation for 20 minutes. Since α-synuclein is thermostable, it remained soluble and was collected with the supernatant. After cooling down the solution to 4 °C, ammonium sulphate was added. For untagged protein, 42 % and for eGFP- and mCherry-tagged, 70 % of saturation was used (tags increased the solubility of α-synuclein in ammonium sulphate solution). Protein precipitation was completed by stirring the mixture at 300 RPM for 30 minutes. After centrifuging the solution at 18 000 x g for 20 minutes, pelleted denatured proteins were dissolved in a buffer solution containing 20 mM Tris-HCl, 0.5 mM DTT and 1 mM EDTA (pH 8.0), filtered and dialyzed for 3 hours in the same buffer solution to remove any excess of ammonium sulphate. After the dialysis was completed, the solution was mixed with DEAE sepharose sorbent and stirred for 30 minutes at 4 °C. The final mixture was loaded onto a HiScale™ 26/20 column for ion-exchange chromatography. α-syn was eluted by increasing the NaCl concentration to 100 mM. Collected fractions containing α-synuclein were concentrated (up to ∼10 mg/mL protein concentration) and loaded onto a HiLoad 26/600 size exclusion chromatography column packed with Superdex 75 resin. The protein was eluted at 2.5 ml/min using PBS. After the procedure was finished, all collected α-syn fractions were mixed together, concentrated and filtered through 0.22 µm mixed-cellulose syringe filters, yielding a final concentration of 600 µM. The protein solution was distributed into fractions of 500 µL and stored at -20°C.

### Fluorescein labelling

Protein fluorescent labelling with fluorescein isothiocyanate (FITC) was done using a modified version of a previously described fluorochrome conjugation protocol ^32^. The α-syn stock solution (100 µL) was combined with 400 µL 100 mM sodium carbonate buffer solution (pH 9.0) and concentrated to 100 µL using a 10 kDa 0.5 mL Pierce protein concentrator (Thermofisher Scientific, cat. No. 88513). These dilution and concentration steps were repeated three times. After the final step, the protein solution was diluted to 1 mg/mL using the sodium carbonate buffer solution. FITC was dissolved in dimethyl sulfoxide under dark conditions to a final concentration of 1 mg/mL. The prepared protein solution (450 µL) was combined with FITC (50 µL) and incubated at 4°C under dark conditions for 24 hours. The solution was then concentrated as described previously to 100 µL and diluted to 500 µL using PBS (pH 7.4). These concentration and dilution steps were repeated 3 times. After the final step, the protein solution was diluted to 400 µM, divided into aliquots of 10 µL and stored at -20°C under dark conditions.

### Liquid-liquid phase separation (LLPS)

To induce protein LLPS, the regular and fluorescently tagged α-syn were placed in an environment containing a crowding agent - polyethylene glycol (PEG, 20 kDa ^33^). PEG was dissolved in MilliQ H_2_O and supplemented with 10x PBS to yield a solution containing 35% PEG (w/v) and 1x PBS (pH 7.4). The α-syn, PEG, PBS and tagged-protein solutions were combined to final reaction mixtures containing 20% PEG, 1x PBS and 200 µM total protein concentration with varying concentrations of tagged proteins. For control samples, tagged proteins were combined with the aforementioned solutions without the addition of regular α-syn. Each sample was incubated for 10 minutes at 22°C in non-binding 1.5 mL test tubes (400 µL final volume) before further examination procedures.

### Light scattering assay

200 µL aliquots of each sample were placed in 3 mm pathlength quartz cuvettes and their right-angle light scattering was scanned using a Varian Cary Eclipse fluorimeter (800 nm excitation and emission wavelengths, 2.5 second signal averaging time). For each condition, three measurements were taken and averaged.

### Fluorescence microscopy

15 μL aliquots of each sample were pipetted onto 1 mm glass slides (Fisher Scientific, cat. No. 11572203), covered with 0.18 mm coverslips (Fisher Scientific, cat. No. 17244914) and imaged immediately after preparation. Images were acquired using Olympus IX83 microscope with 40x objective and fluorescence filter cubes (480 nm excitation and 530 nm emission wavelengths for eGFP- and FITC-α-syn, 545 nm excitation and 600 nm emission wavelengths for mCherry-α-syn). Twelve different images were taken from each condition and analyzed using ImageJ software ^34^. Identical background subtraction and contrast/brightness settings were applied to all images.

### Electron microscopy

eGFP- and mCherry-α-syn samples were prepared 5 minutes before their application to TEM grids. Five microliters of each 100 μM alpha-synuclein aggregate solution was applied to the glow-discharged formvar/carbon supported 300 mesh copper grids (Agar Scientific, U.K.) for 1 minute. After the excess fluid was removed with filter paper, the grid was washed once with 5 μL of water for 1 minute to remove PEG residues before staining the sample. Next, the grid was negatively stained with 5 μL of 2% (w/v) uranyl acetate for 1 minute, followed by five 1 minute washes with 5 μL of water. All TEM images were acquired on a Talos 120C (Thermo Fisher) microscope operating at 120 kV and equipped with a 4k × 4k Ceta CMOS Camera and after that were analyzed using ImageJ software.

### Seeded aggregation

Samples containing 20 µM of mCherry- or eGFP-α-syn and 20% PEG were incubated at 22°C for 1 hour, after which they were centrifuged at 14 000 x g for 15 min. The supernatants were then removed and the pelleted aggregates were resuspended into an identical volume of PBS. This centrifugation and resuspension procedure was repeated 3 times. The resulting solutions were then combined with PBS, 10 mM thioflavin-T (ThT) and α-syn stock solutions to a final monomeric protein concentration of 100 µM, 100 µM ThT and 5% (v/v) of the aggregate solution. The control sample contained PBS in place of the aggregate solution. The reaction solutions were distributed to 96-well plates and aggregation was monitored using a ClarioStar Plus plate reader under constant 600 RPM agitation and 37°C temperature. Measurements were taken every 5 min, using 440 nm excitation and 480 nm emission wavelengths. The resulting kinetic data was fit using a Boltzmann sigmoidal equation using Origin 2018 software. The reaction lag time and apparent rate constants were determined as described previously ^35^.

### ThT binding assay

mCherry- and eGFP-α-syn samples were prepared, incubated and pelleted as described in the seeded aggregation method section. The supernatants were removed and the pellets were resuspended into identical volumes of PBS, which contained 100 µM ThT. The samples were then placed into 3 mm pathlength quartz cuvettes and scanned using a Varian Cary Eclipse fluorimeter, using a 440 nm excitation wavelength and measuring the fluorescence between 460 and 520 nm (1 nm steps).

### Denaturation assay

Due to the high stability of mCherry and eGFP fluorescent tags ^36^, different concentrations of each tagged protein were resuspended into PBS (pH 7.4) containing 4 M guanidinium hydrochloride – a strong denaturing agent ^37^. Thermal denaturation of mCherry- and eGFP-α-syn was measured by tracking the fluorescence intensity of each protein using a Varian Cary Eclipse fluorimeter. For each protein type and concentration, three separate 200 µL samples were placed in 3 mm pathlength cuvettes and incubated at 20°C for 2 minutes. Temperature was increased from 20°C to 98°C at a rate of 2°C/min. Sample fluorescence intensity was scanned every 30 seconds (2.5 second signal averaging time, 488 nm excitation – 510 nm emission wavelengths for eGFP and 585 nm excitation – 610 nm emission wavelengths for mCherry).

Protein denaturation midpoint temperature (T_m_) was determined as the main minimum of the first derivative of the thermal denaturation profile, assuming the loss in fluorescence intensity stems from the disassembly of the mCherry and eGFP protein structures.

### Size exclusion chromatography

The size of alpha-synuclein and its fluorescently tagged variants was assessed by size exclusion chromatography (Tricorn 10/300 column (Cytiva)), packed with Superdex 75 prep grade resin (Cytiva). The column was precalibrated with PBS and 250 μL of 50 μM of selected protein solution was injected using ÄKTA Start (Cytiva) chromatography system.

### Droplet statistical analysis

For each condition, twelve fluorescence microscopy images were used to determine the size and distribution of α-syn droplets. Droplet parameters were analyzed with ImageJ software ^34^ using automatic threshold selection and particle analysis. For further statistical analysis, all particles of 4 or less pixel size were regarded as artefacts and not taken into account. The total droplet count was the sum of all particles detected in all twelve images for each condition. Droplet volume was calculated based on the particle areas (assuming completely spherical condensates).

## Results

For an initial assessment of tagged α-syn influence on the process of LLPS, different concentrations of mCherry-, eGFP- and FITC-α-syn samples were examined using right-angle light scattering (Figure 1). When the tagged proteins were in solution without regular α-syn (Figure 1A), no effects on sample turbidity were observed up to a concentration of 1 µM. Past this point, the light scattering data diverged, with the 2 µM eGFP-α-syn sample having a significantly higher signal value (ANOVA One-way Bonferroni means comparison, n=3, p<0.05). At 4 µM, the mCherry-α-syn sample also showed signs of turbidity changes, while eGFP-α-syn displayed an even higher scattering intensity. This concentration-dependent increase in samples light scattering persisted for both the mCherry-α-syn and eGFP-α-syn samples, with both protein samples having significantly higher intensity values than the control at the largest tested concentration. Conversely, FITC-α-syn demonstrated an opposite effect, where even the 20 µM sample did not deviate from the baseline value and was significantly lower than the control.

**Figure 1.**
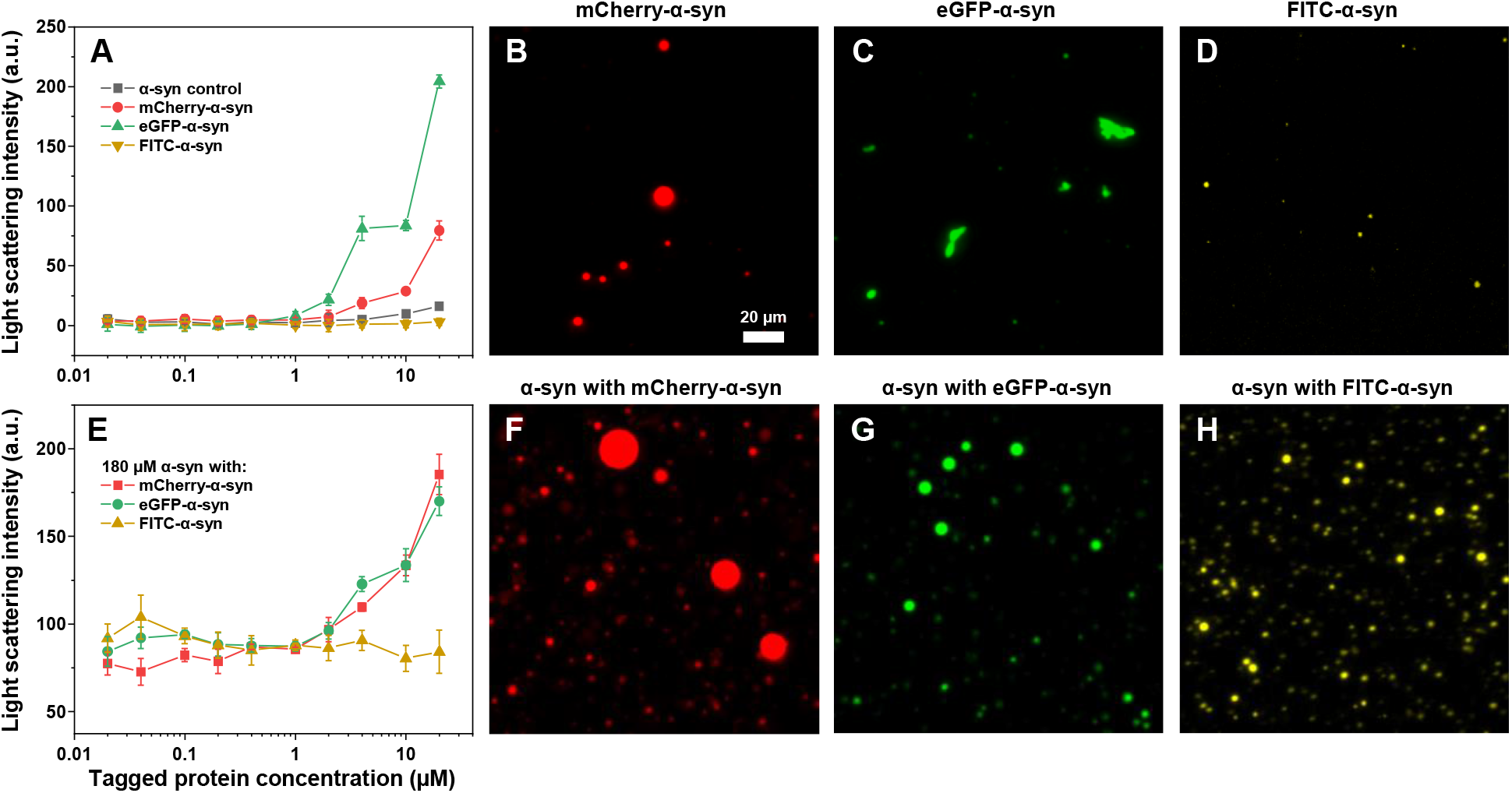
Influence of fluorescent protein tags on α-syn droplet and aggregate assembly. Right-angle light scattering of different concentration mCherry-, eGFP- and FITC-α-syn in the absence (A) or presence (E) of regular α-syn. Fluorescence microscopy images of 20 µM mCherry-, eGFP- and FITC-α-syn in the absence (B – D) and presence (F – H) of regular α-syn. Sample measurements were taken 10 minutes after their preparation. For each condition, three light scattering measurements were taken and averaged, error bars are for one standard deviation. For fluorescence microscopy, 12 images were taken under each condition (scale bar shown in panel B (20 µm) is identical for all images, additional images available as Supporting Information).

Examining the 20 µM tagged protein samples with fluorescence microscopy revealed that each group exhibited different types of self-association. In the case of mCherry-α-syn, the samples contained protein droplets, ranging in size from less than 1 µm to over 10 µm in diameter (Figure 1B). eGFP-α-syn, on the other hand, assembled into either amorphous or elongated aggregates (Figure 1C, Supporting Figure S1), which would explain their relatively higher level of light scattering among all the tested groups. FITC-α-syn formed a relatively small number of tiny assemblies (Figure 1D), most of which were only faintly visible during the fluorescence microscopy assay. In this case, the observed structures could not be categorized as droplets or protein aggregates and may be microscopy artefacts. The structures formed by mCherry- and eGFP-α-syn were further examined using electron transmission microscopy. The images did not reveal any clear signs of amyloid-like fibrils and the samples contained mostly amorphous structures (Supporting Figure S2).

To determine how these tagged proteins influenced LLPS of regular α-syn, it was combined with different ratios of mCherry-, eGFP- and FITC-α-syn to a final protein concentration of 200 µM. When the samples were examined using right-angle light scattering (Figure 1E), no notable deviations from their respective controls were observed up to a concentration of 4 µM, apart from small stochastic fluctuations in scattering intensity (0.04 µM and 0.1 µM conditions). Unlike the previous examination with only tagged proteins, the mCherry- and eGFP-α-syn samples displayed similar concentration-dependent changes in scattering intensity, with both the 10 µM and 20 µM condition values being within margin of error. FITC- α-syn retained a similar tendency as was observed in the previous assay with no noteworthy deviations through the entire concentration range and the 20 µM sample value was not significantly different from the control.

During fluorescence microscopy examination of the highest tagged protein concentration samples, all three protein groups resulted in the formation of droplets (Figure 1F-H). For mCherry-α-syn, the samples contained a number of very large protein assemblies, with some exceeding a diameter of 20 µm (Figure 1F). Unlike the sample without regular α-syn, there was also a large quantity of smaller droplets alongside their massive counterparts. In the case of eGFP-α-syn, the presence of regular α-syn prevented the formation of eGFP-α-syn aggregates and the proteins underwent LLPS (Figure 1G). The formed droplets were noticeably smaller than the mCherry-α-syn variants and they appeared to be more abundant in quantity. FITC-α-syn samples contained the smallest droplets out of the three groups (Figure 1H), as well as the presence of a vast number of tiny, faintly visible assemblies, mirroring the results of the previous examination.

To gain a deeper insight into this tagged protein-induced change to α-syn LLPS, 200 µM protein concentration samples, containing a range of mCherry-, eGFP- and FITC-α-syn were examined using fluorescence microscopy. In the case of mCherry-α-syn, samples with 1 µM (0.5%), 2 µM (1%) and 4 µM (2%) of the tagged protein did not display any visually distinct features (Figure 2A). When the concentration was 10 µM (5%), there was a notable increase in the number of larger size droplets. This effect was further accentuated at 20 µM (10%) mCherry-α-syn, where the vast majority of the protein was accumulated in large 10 – 30 µm diameter droplets.

**Figure 2.**
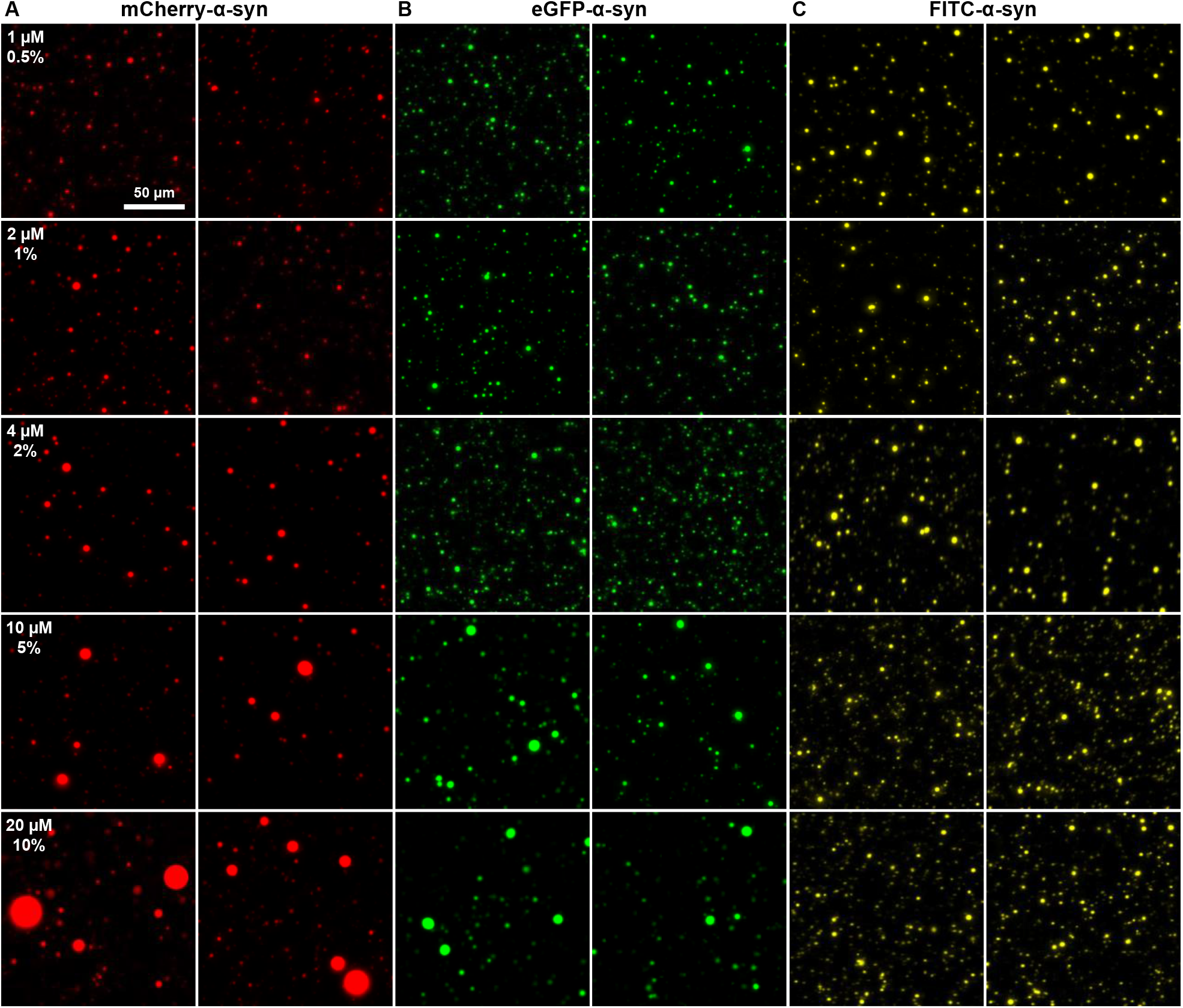
Fluorescence microscopy images of α-syn samples containing different fluorescent tags. Representative images of regular α-syn combined with different concentrations of mCherry- (A), eGFP- (B) and FITC-α-syn (C) to a final concentration of 200 µM. For each condition, twelve images were obtained (available as Supporting Information). Scale bar (50 µm) shown in the top left panel is identical for all images.

When the protein mixture contained eGFP-α-syn, a similar event of larger condensate formation was observed for the 10 µM and 20 µM concentration samples (Figure 2B). However, the droplets were significantly smaller in size when compared to the mCherry-α-syn variant. Interestingly, there appeared to be a gradual increase in the number of tiny protein condensates, with the largest number detected at 4 µM eGFP-α-syn (Supporting Figure S3). At higher concentrations, there was a notable decrease in such small α-syn accumulations, which coincided with the appearance of larger droplets.

When the same examination was conducted with FITC-α-syn, there were no clear signs of larger droplet formation as a function of the tagged protein concentration (Figure 2C). Similar to eGFP-α-syn, there was a concentration-dependent increase in the number of tiny protein condensates (Supporting Figure S3), albeit without a discontinuity marked with a change in droplet size. These results, combined with the two previous tagged protein groups suggested that each fluorescent tag had a distinct effect on the droplet formation process.

In order to quantify the influence of tagged proteins on α-syn LLPS and to determine the concentration thresholds for each type of alteration to condensate formation, an analysis was performed using the larger array of sample images (available as Supporting Information). Droplets were divided into five groups based on their volume to evaluate which group was dominant under each condition.

When regular α-syn samples contained small concentrations of mCherry-α-syn, most of the protein was condensed into 10 – 100 µm^3^ volume droplets, followed second by 100 – 1000 µm^3^ particles (Figure 3, red). At 4 µM mCherry-α-syn, the distribution shifted, and majority of the protein was contained within 100 – 1000 µm^3^ condensates. At even higher tagged protein concentrations (10 µM and 20 µM), most of α-syn was found within droplets larger than 1000 µm^3^ in volume. When only the number of condensates were compared, all conditions displayed that 1 – 10 µm^3^ volume assemblies were the dominant type.

**Figure 3.**
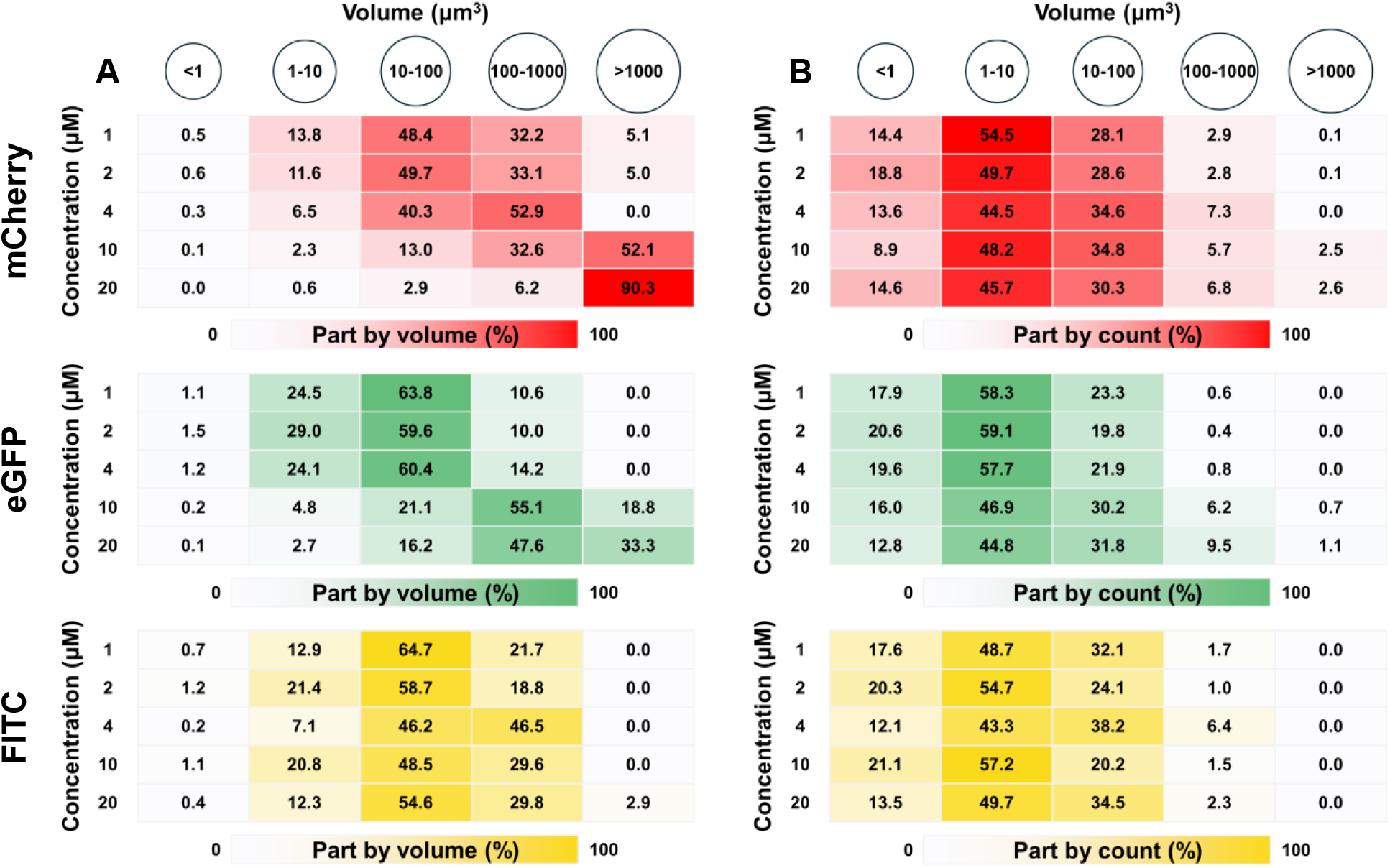
Differently tagged α-syn influence on the number and volume of droplets. Distribution of droplets by volume (A) and count (B) in samples containing mCherry-, eGFP- and FITC-α-syn (12 images for each condition, all particles above 4 pixels in size).

In the case of eGFP-α-syn, a similar trend was observed at low tagged protein concentrations, with most of the protein assembling into 10 – 100 µm^3^ droplets, however, unlike with mCherry-α-syn, the second largest group was 1 – 10 µm^3^ condensates (Figure 3, green). Another clear difference was the higher concentration of eGFP-α-syn required for a transition to higher volume droplets to occur (10 µM as opposed to 4 µM). Complementing previously observed results, the highest concentration of eGFP-α-syn did not cause α-syn to condense into droplets larger than 1000 µm^3^ in volume and the 100 – 1000 µm^3^ particles remained the dominant type. Based on the count of eGFP-α-syn droplets, the 1 – 10 µm^3^ condensates were the most abundant, similar to the mCherry-α-syn variant.

As expected, the FITC-α-syn group of samples did not follow a concentration-dependent trend as both previous fluorescent tags and the majority of α-syn was condensed into 10 – 100 µm^3^ volume droplets (Figure 3, yellow). In the case of 4 µM FITC-α-syn, however, there appeared to be a shift towards larger droplets, as seen by the nearly equivalent distribution between 10 – 100 and 100 – 1000 µm^3^ volume groups. Based on the count of droplets, no notable deviations were observed for any concentration and the distribution was similar to mCherry- and eGFP-α-syn.

Since both mCherry- and eGFP-α-syn variants were capable of droplet/aggregate formation under LLPS-inducing conditions and modulated α-syn condensation, it was hypothesized that this effect may stem from their ability to associate in solution. To examine this possibility, the two tagged proteins were combined with regular α-syn and analyzed using size exclusion chromatography (SEC). When the proteins were separate, mCherry- and eGFP-α-syn had almost identical elution profiles (Figure 4A). After they were combined with regular α-syn, there were minimal changes to the elution profile peak positions (Figure 4B), indicating that the proteins did not form significant cross-interactions.

**Figure 4.**
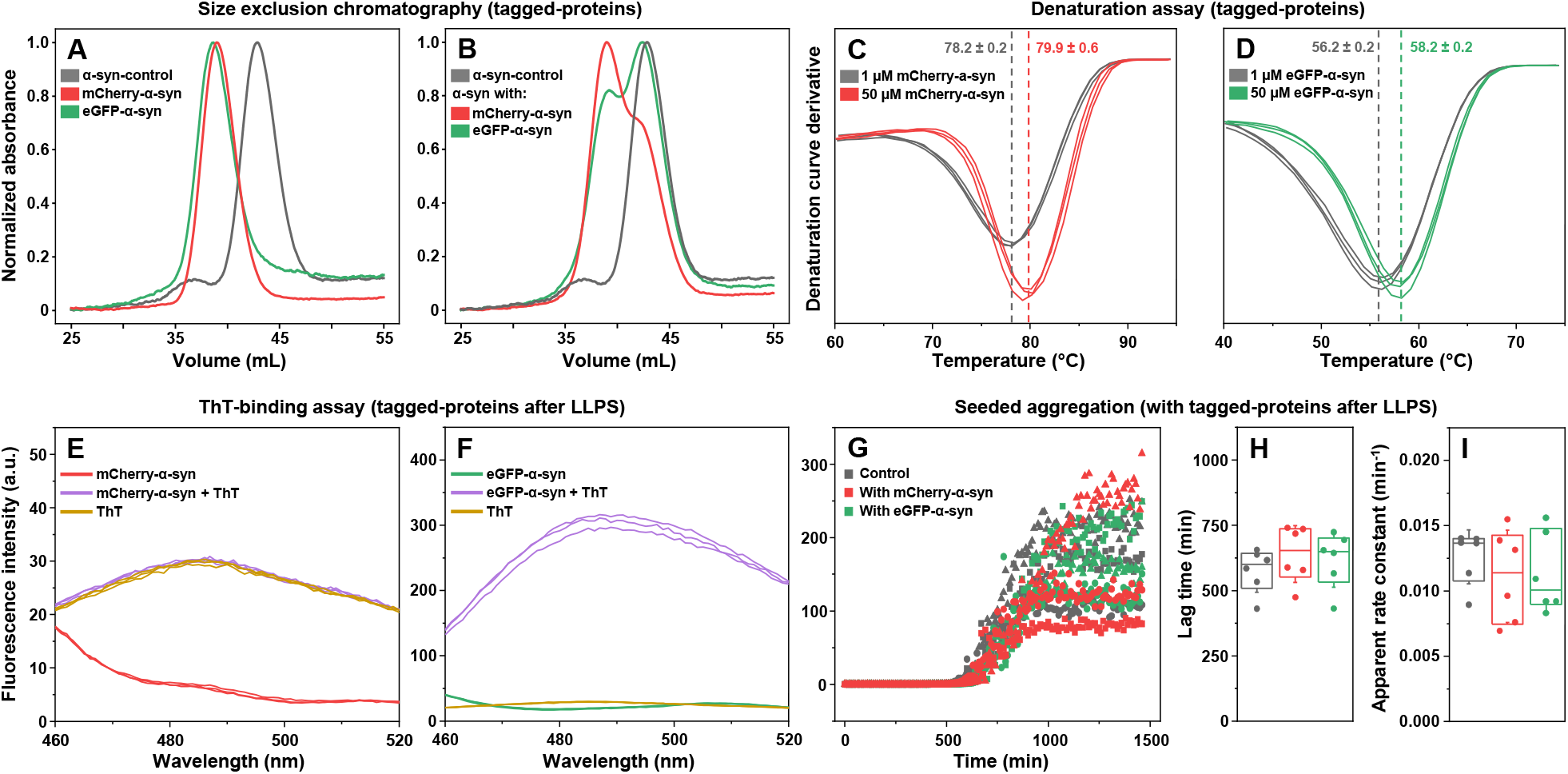
Examination of fluorescently tagged α-syn cross-interaction and amyloid-like characteristics. Size exclusion chromatograms of 50 µM mCherry-α-syn and eGFP-α-syn when in the absence (A) or presence (B) of 50 µM regular α-syn. Thermal denaturation curve derivatives of different concentration mCherry-α- syn (C) and eGFP-α-syn (D). ThT-binding assay of mCherry- (E) and eGFP-α-syn (F) sample aggregates after resuspension into PBS. Seeded aggregation kinetics (G), reaction lag time (H) and apparent rate constant (I) distribution of regular α-syn with the addition of mCherry- or eGFP-α-syn aggregates.

Another probable event was a concentration-dependent self-association of the tagged proteins. To examine this possibility, thermal denaturation assays were performed on different concentration samples of mCherry- and eGFP-α-syn, in the presence of 4 M guanidinium hydrochloride (Figure 4C, D). When comparing 1 µM and 50 µM samples, it was observed that the higher concentration of both proteins resulted in an approximately 2 degree increase in their T_m_ values. These results suggested that there may be a concentration-dependent change in their cross-interaction, leading to a small increase in the protein structural stability. Interestingly, despite both proteins having only minor distinctions in sequence and structure, the difference in their T_m_ values was more than 20 degrees, with mCherry-α-syn being the more stable structure. The higher stability of mCherry-α-syn correlated with the formation of significantly larger droplets under LLPS-inducing conditions.

Another aspect that required further investigation was the tagged protein self-assembly in the absence of regular α-syn. While mCherry-α-syn was able to form large droplets at relatively low concentrations, eGFP-α-syn associated into aggregates, which could potentially be amyloid-like structures. To examine this possibility, the 20 µM tagged protein samples under LLPS conditions were centrifuged and the pelleted structures were resuspended into PBS without the crowding agent. Interestingly, both mCherry- and eGFP- α-syn samples contained small amounts of insoluble structures. The samples were then combined with an amyloid-specific dye – thioflavin-T (ThT) and their fluorescence spectra were acquired. In the case of mCherry-α-syn (Figure 4E), the spectra were nearly identical to the control sample, indicating that the insoluble particles did not bind the fluorescent dye. The eGFP-α-syn sample, however, displayed a 10-fold higher fluorescence intensity than the control (Figure 4F), suggesting that it formed structures capable of binding ThT. Whether these assemblies were amyloid structures or could simply bind ThT in a non-specific manner and enhance its fluorescence intensity ^38^ remained questionable.

To determine if the insoluble particles were amyloid-like, a seeded aggregation experiment was conducted. The mCherry- and eGFP-α-syn samples were combined with regular α-syn in a 1:20 ratio and their aggregation reactions were monitored (Figure 4G). Surprisingly, neither tagged protein had any effect on the aggregation reactions, with no significant differences being determined between their lag times (Figure 4H) or apparent rate constants (Figure 4I, n=6, p<0.05). These results indicate that the structures formed by the two tagged proteins are likely amorphous aggregates, which are incapable of affecting the amyloid fibril formation of regular α-syn.

## Discussion

Over the last decade, the process of LLPS, its role in various cellular functions, and implication in the onset of several widespread disorders has garnered significant attention from the scientific community. In the process of studying biomolecule condensate formation, a variety of fluorescent probes have been applied, including different color protein-based tags, as well as covalently bound chemical compounds ^8,12,14,16^. Despite the common application of such fluorescent labels, very little is known regarding their influence on the process of LLPS, with only a handful of reports providing examples of their potential downsides ^19,21,22^. In this work, we examined the influence of three fluorescent tags on the process of α-syn phase separation and discovered that each protein modification had unique, LLPS-altering properties.

The most notable and clearly observable effect of fluorescent labels was the change in α-syn droplet size. While the lowest tested concentrations of tagged proteins did not display any significant changes, droplet sizes were influenced by the presence of 4 µM and higher concentrations of mCherry- and eGFP-tagged proteins. This specific concentration was also the only condition where FITC-α-syn had a noteworthy effect on the condensate size distribution. Taking into consideration the range of different fluorescent label concentrations used *in vitro* ^14,15,23,24^, as well as lack of complete control over their distribution during *in vivo* LLPS studies, it is important to take the tag-induced modulation of condensate formation into account.

Apart from their influence on droplet size, larger tagged protein concentrations also modulated the overall number of detectable particles over the same area of fluorescence microscopy images (Supporting Figure S3). This effect was unique for all three tested fluorescent labels, with the highest droplet count observed in the 4 µM eGFP-α-syn samples. While FITC-α-syn displayed a concentration-dependent trend of increasing particle count and mCherry-α-syn reduced the number of condensates due to the formation of massive droplets, eGFP-α-syn had a point of discontinuity at the 4 µM tagged protein concentration. Under this condition, the particle count was more than two times higher than under both the lower and higher concentration. This was also the point which marked the shift from 10 – 100 µm^3^ to 100 – 1000 µm^3^ volume particle dominance. These results suggest that even small alterations in fluorescently labeled proteins can have drastic effects on the process of α-syn LLPS.

Another interesting aspect was the self-association tendencies of mCherry- and eGFP-α-syn in the absence of regular α-syn. Under the tested conditions, FITC-α-syn did not associate into detectable droplets (Figure 1D), while both aforementioned protein-based fluorescent labels induced the formation of either predominantly large droplets (mCherry, Figure 1B) or aggregates (eGFP, Figure 1C). These results both complement previous observations that proteins tags can promote the process of LLPS ^19,20^, as well as raise concerns regarding the sample preparation methodologies for *in vitro* studies. This is especially evident in the case of eGFP-α-syn, where a mixture of regular α-syn and the tagged protein resulted in only droplet formation, while eGFP-α-syn alone associated into amorphous structures which could bind an amyloid-specific dye molecule, without affecting the aggregation of regular α-syn. These findings suggest that the order of component mixing is critically important when using fluorescently tagged proteins for *in vitro* experimental procedures.

## Conclusions

Based on the results of this study, it is clear that fluorescently labeled proteins introduce the problem of altering the characteristics of condensates during LLPS. Since this effect of droplet formation is concentration dependent, it is advisable to use the lowest possible tagged protein concentration, which still facilitates the ability to detect LLPS via fluorescence microscopy. The labeled proteins should also be placed in the sample as the final component, in order to prevent their self-association into droplets or amorphous aggregate structures.

## Supporting information

Images and raw data

## Data availability

All raw data and additional fluorescence microscopy images are included as Supporting information.

