## Supplementary information for "Diverse effects of fluorescent labels on alpha-synuclein condensate formation during liquid-liquid phase separation"

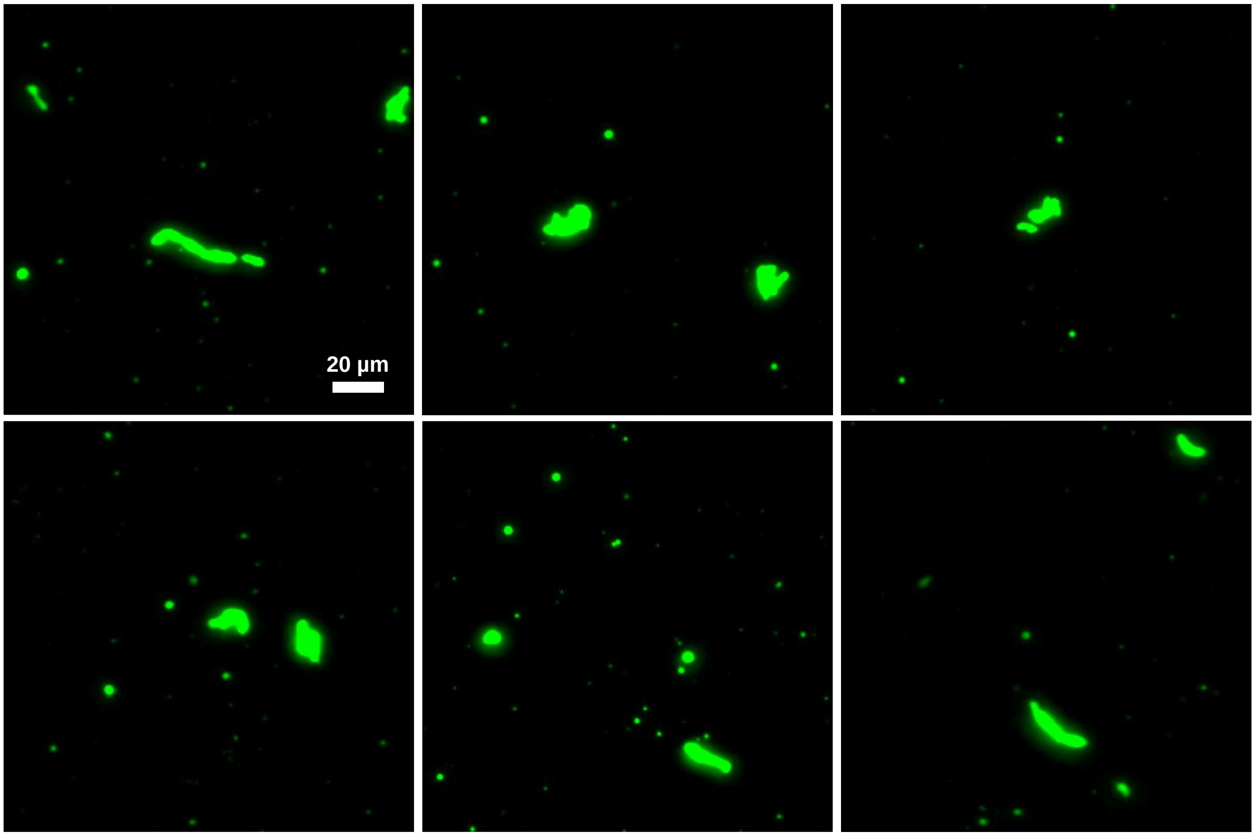


**Figure S1.** Fluorescence microscopy images of eGFP-α-syn aggregates formed under LLPs conditions in the absence of regular α-syn (scale bar = 20 μm).

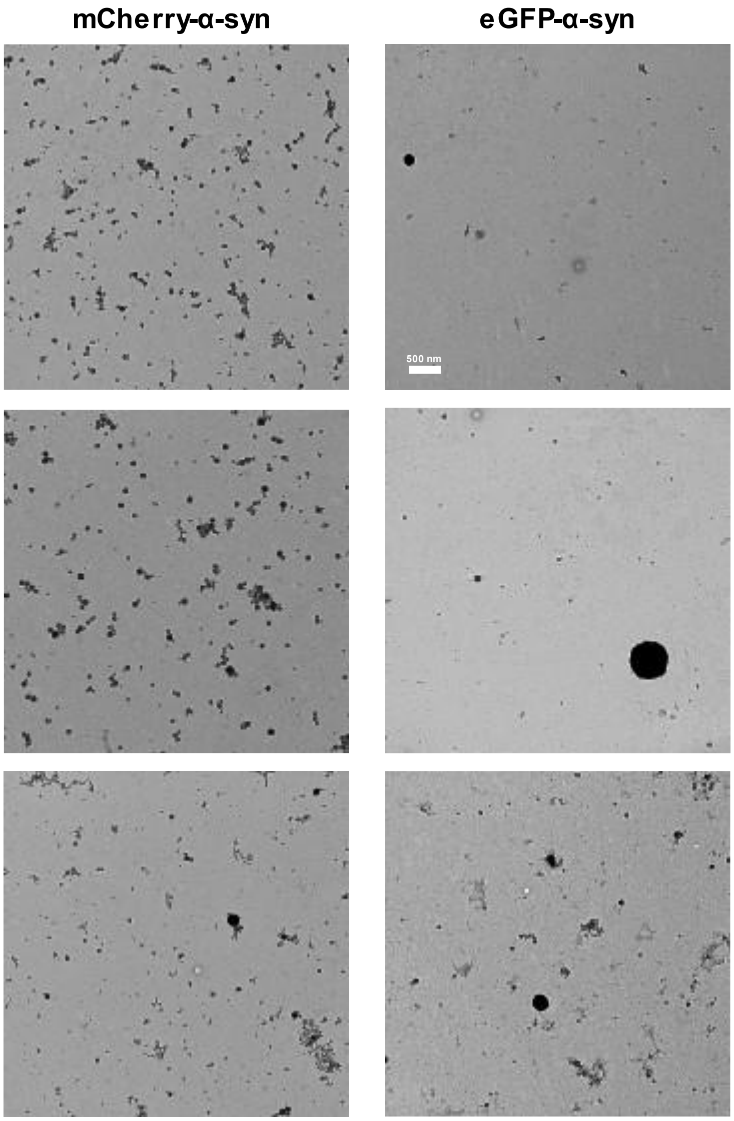


**Figure S2.** Transmission electron microscopy images of mCherry- and eGFP-α-syn structures, formed in the absence of regular α-syn under LLPS-inducing conditions (scale bar = 500 nm).

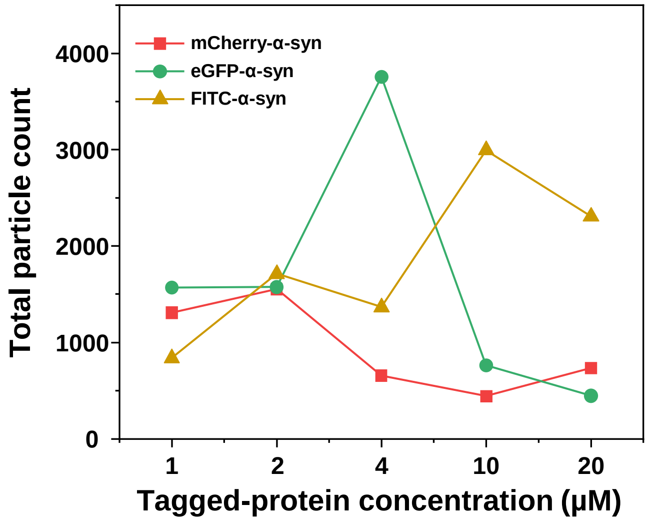


**Figure S3.** Total particle count under different tagged-protein concentrations (all particles above 4 pixels in size from 12 images under each condition).
